# TSNAdb v2.0: the updated version of tumor-specific neoantigen database

**DOI:** 10.1101/2022.07.28.501872

**Authors:** Jingcheng Wu, Wenfan Chen, Yuxuan Zhou, Ying Chi, Xiansheng Hua, Jian Wu, Xun Gu, Shuqing Chen, Zhan Zhou

**Author notes:** To whom correspondence should be addressed. Corresponding author. (Zhou Z).

## Abstract

Tumor neoantigens have been well-acknowledged as ideal targets for tumor immunotherapy in recent years. With the deepening of research on neoantigen-based tumor immunotherapy, comprehensive neoantigen databases are urgently needed to meet the growing demand for clinical studies. We have built the Tumor-Specific NeoAntigen Database (TSNAdb v1.0) previously, which has attracted wide attention. In this study, we provide an updated version of the Tumor-Specific NeoAntigen Database (TSNAdb v2.0) with several new features including 1) taking stricter criteria for neoantigen identification. 2) providing predicted neoantigens derived from three types of somatic mutations. 3) collecting experimentally validated neoantigens and classifying them according to the evidence. TSNAdb v2.0 is freely available at https://pgx.zju.edu.cn/tsnadb/.

## INTRODUCTION

Tumor neoantigens are tumor-specific antigens derived from somatic mutations in tumor cells, which have been well-acknowledged as ideal targets for tumor immunotherapy in recent years [1–4]. Because of the large workload of experimental verification, the practical solution for neoantigen identification is to take advantage of cancer genomics and bioinformatics. Numerous prediction tools considering the biological process of neoantigen generation such as human leukocyte antigen (HLA)-peptide binding [5–7] have been developed, which have been embedded in neoantigen prediction pipelines such as pVACtools [8], TSNAD [9,10], and pTuneos [11]. Neoantigen-related databases such as TCLP [12], TCIA [13], and TSNAdb [14] also have been created for the better usage of neoantigen in clinical research. In the first version of TSNAdb, we took the complex of mutated peptides and the HLA class I molecules (peptide-HLA pairs, pHLAs) as tumor neoantigens and predicted binding affinities between mutated/wild-type peptides and HLA class I molecules by NetMHCpan v2.8/v4.0. 3,707,562/1,146,961 potential neoantigens derived from single nucleotide variants (SNVs) of 7,748 tumor samples from The Cancer Genome Atlas (TCGA) were presented in the database. With the gradual deepening of research on neoantigen-based tumor immunotherapy, neoantigens from other types of mutations have been identified and more experimental data have been generated. So it is urgent to perform system updates for the tumor-specific neoantigen database.

We present here a major updated version of TSNAdb that implements new features and improvements including 1) Stricter criteria are used for neoantigen identification to reduce the high false-positive rate of neoantigen prediction in practice. Only the pHLAs that meet the thresholds of three tools (DeepHLApan [5], MHCflurry [6], and NetMHCpan v4.0 [7]) would be considered as potential neoantigens. The HLA-peptide pairs would also not be considered neoantigens if the mutated genes are not expressed in the tumor cells. 2) Providing the predicted neoantigens derived not only from SNVs but also from insertions/deletions (INDELs) and gene fusions (Fusions). Totally, 372,273 SNV-derived neoantigens, 137,130 INDEL-derived neoantigens and 11,093 Fusion-derived neoantigens are obtained. The mean number of neoantigens each SNV (0.38) generates is lower than that each INDEL (1.22) or Fusion (0.88) generates. 3) Collecting as much as experimentally validated neoantigens from public databases and literature. 1,856 neoantigens are collected and divided into three tiers due to the level of experimental verification. Corresponding genes and mutations of the collected neoantigens are linked to CandrisDB (Cancer-driving site profiling database) [15], providing more convincible information since shared neoantigens derived from driver genes or driver mutations would be ideal targets for tumor immunotherapy [16].

We believe that the updated database would contribute to the neoantigen-based tumor immunotherapy and the database would continue updating in the aspect of predicting neoantigens from more types of mutations and collecting more experimentally validated neoantigens.

## MATERIALS AND METHODS

### Data collection and preprocessing

The SNVs, INDELs, and the expression level of corresponding genes are collected from TCGA (https://portal.gdc.cancer.gov/). Mutated nucleotide sequences generated by SNVs and INDELs are translated into mutated amino acid sequences and have been decomposed into 8 to 11 peptides using the pipeline of TSNAD v2.0[10]. The gene fusions are collected from Gao et.al [17] and the mutated proteins are generated by STAR-FUSION [18] as the literature stated. The HLA alleles of corresponding samples are collected from TCIA (https://tcia.at/home). Finally, 972,187 SNVs from 7,748 samples, 112,404 INDELs from 7,086 samples and 12,639 Fusions from 4,234 samples were used for neoantigen prediction.

### Neoantigen prediction

Three tools (DeepHLApan, MHCflurry, and NetMHCpan v4.0) are used for neoantigen prediction and only the pHLAs that meet all the criteria of the three tools would be considered potential neoantigens. The reason we choose these three tools are as follows: NetMHCpan is the most frequently used tool for neoantigen prediction in clinical practice. MHCflurry obtains the prediction neoantigen efficiently and with high quality. DeepHLApan considers both HLA-peptide binding and immunogenicity of pMHC that the other two tools haven’t taken into consideration for high-confidence neoantigen prediction. In this study, we select the threshold of each tool as follows. For NetMHCpan 4.0, pMHC with rank < 2% or affinity <500 nM is considered binding, we take both thresholds to select higher quality neoantigen. The output of MHCflurry is rank% and has no specific threshold, we set rank < 2% as the threshold which is the same as NetMHCpan 4.0. The predicted scores of DeepHLApan are posterior probability so we set the threshold to 0.5. Besides, the pHLAs whose corresponding genes are not expressed (transcripts per million reads, TPM < 1) are removed.

### Experimentally validated neoantigens

The validated neoantigens were collected not only from several neoantigen databases (dbPepNeo [19,20], NeoPeptide [21], NEPdb [22], CAPD [23]) but also from published literature through data mining. For the neoantigens without gene or mutation information, BLAST was used to figure out the mutated genes and the positions of somatic mutations at proteins. All collected data would be further checked whether the neoantigens are immunogenic or being presented to the cell surface, and the collected neoantigens are divided into three tiers according to the experimental level. Neoantigens which have been both validated as immunogenic and to be presented to the cell surface are labeled tier1, while those only validated as immunogenic are labeled tier2, and those only validated to be presented to the cell surface are labeled tier3.

## RESULTS

### Stricter criteria for neoantigen identification

Neoantigen-based tumor immunotherapy has shown good application prospects in clinical practice. However, the high false-positive rate of neoantigen prediction limits its usage. And how to select high-confidence immunogenic neoantigens remains to be resolved. To reduce the potential false-positive rate in our predicted results, the expression level of the mutated gene was taken into consideration and three well-acknowledged tools were used for neoantigen prediction. Only the pHLAs satisfied the criteria of all tools are claimed as neoantigens (see Methods in detail). Under the new selection criteria, the final number of predicted neoantigens derived from SNVs is 372,273, almost one-third of the neoantigens provided in TSNAdb v1.0 (the number predicted by NetMHCpan v4.0).

### Neoantigens derived from three types of mutation

Neoantigens are not only generated from SNVs but also generated from other mutations such as IDNELs and Fusions. Based on the analysis of different types of mutations in TCGA tumor samples, we provided 137,130 INDEL-derived neoantigens and 11,093 Fusion-derived neoantigens into TSNAdb v2.0. The predicted neoantigens derived from SNVs are more than that derived from INDELs and Fusions due to the greatest number of SNVs in collected somatic mutations. However, the average number of neoantigens derived from each SNV (0.38) is less than that derived from each INDEL (1.22) or each Fusion (0.88) (Table 1). We further explored the relationship between the number of mutations and neoantigens for three mutation types. The results show that the number of SNV-derived neoantigens and INDEL-derived neoantigens have positive correlations with the number of SNVs and INDELs, respectively (Figure 2a,2b, r=0.925 and 0.902, respectively). There is no significant correlation between the number of Fusion-derived neoantigens and the number of Fusions (Figure 2c, r=0.452) which might be attributed to the fact that the impact of fusions varies a lot. In some fusion genes, only the peptides around the fusion sites are neo-epitopes (result in a fewer number of predicted neoantigens), while in other fusion genes, all peptides after fusion sites are neo-epitopes (result in a larger number of predicted neoantigens). A sample with a fewer number of fusions with a bigger impact may have bigger numbers of fusion-derived neoantigens compared with a sample with a larger number of fusions with a fewer impact. Besides, the small number of fusions in each sample increases the difficulty of obtaining a clear relationship between the number of Fusion-derived neoantigens and the number of Fusions.

**Figure 1.**
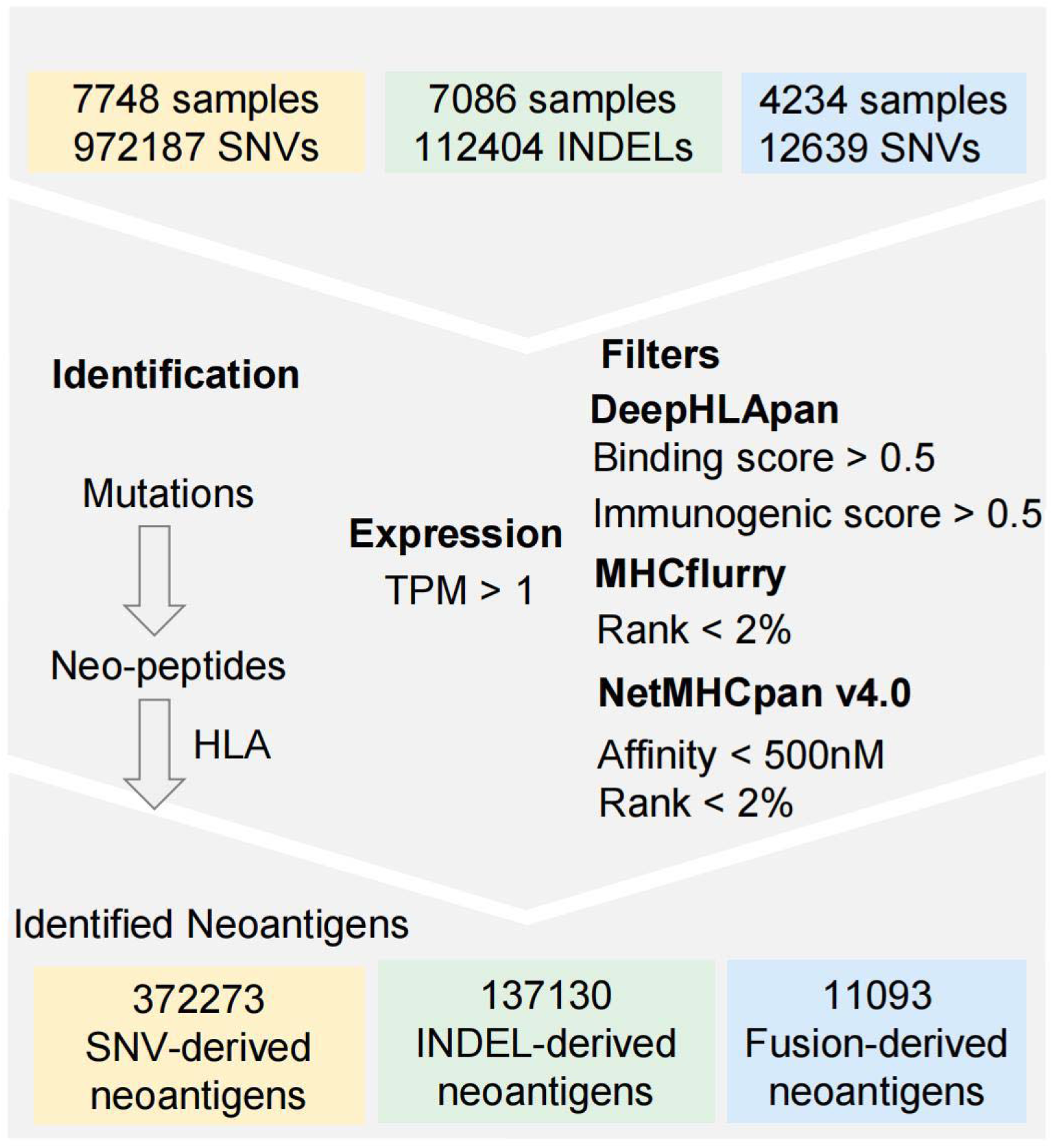
The neoantigen prediction process of TSNAdb v2.0.

**Figure 2.**
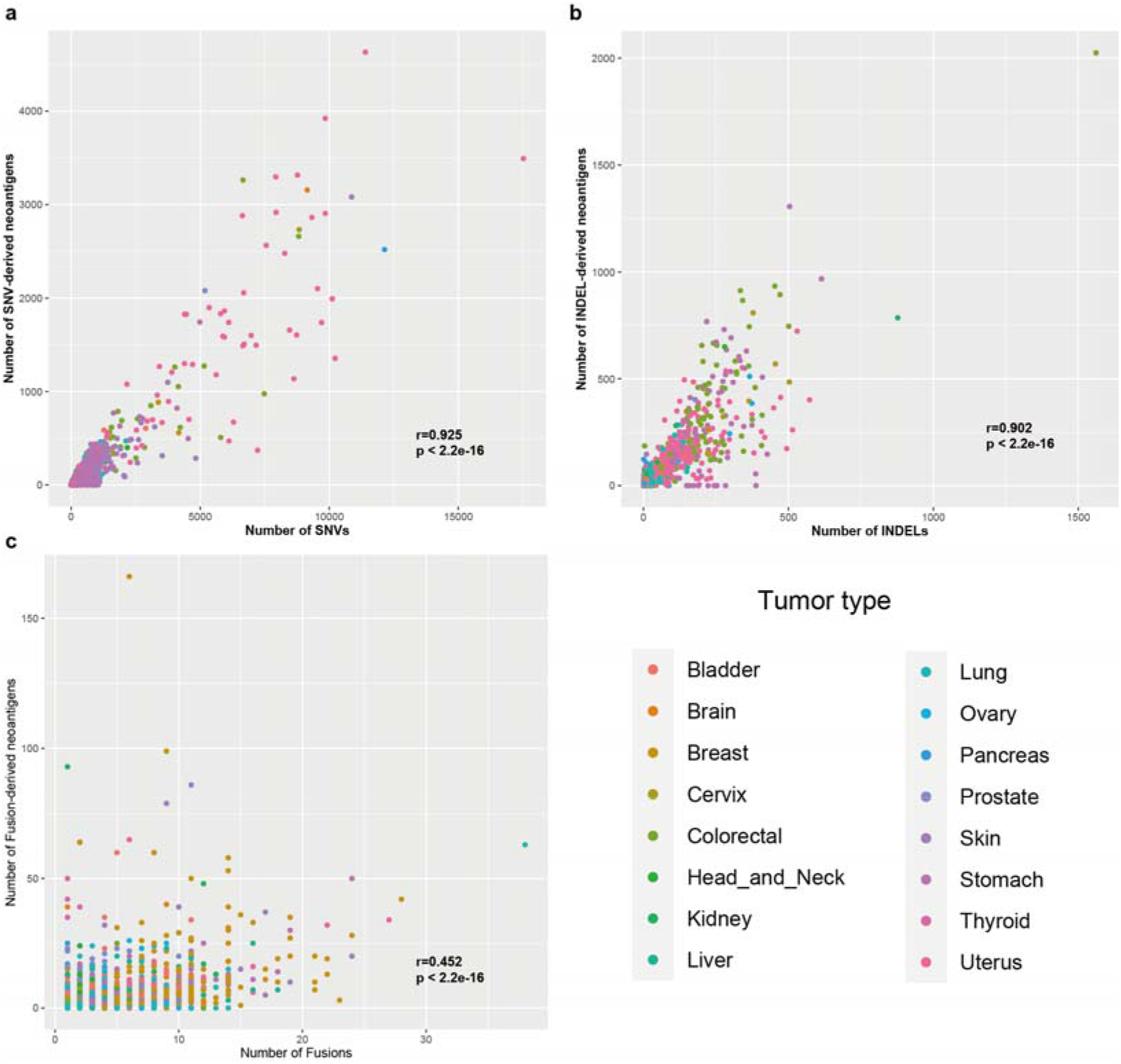
The relationship between the number of mutations and neoantigens across 16 tumor types of three mutation types. **a.** The relationship between the number of SNVs and SNV-derived neoantigens. **b.** The relationship between the number of INDELs and INDEL-derived neoantigens. **c.** The relationship between the number of Fusions and Fusion-derived neoantigens. Pearson correlation coefficient is used for evaluation.

**Table 1.** The distribution of mutations and neoantigens across 16 tumor types.

| Mutation type | Tumor type | No. of samples | No. of mutations | No. of neoantigens | Neoantigen /Mutation |
| --- | --- | --- | --- | --- | --- |
| SNV | Bladder | 408 | 74707 | 21070 | 0.38 |
|  | Brain | 151 | 16197 | 5227 |  |
|  | Breast | 982 | 62211 | 18238 |  |
|  | Cervix | 286 | 46607 | 11906 |  |
|  | Colorectal | 531 | 165293 | 39116 |  |
|  | Head and Neck | 495 | 55968 | 13728 |  |
|  | Kidney | 681 | 26170 | 8499 |  |
|  | Liver | 358 | 28024 | 5987 |  |
|  | Lung | 999 | 212861 | 52801 |  |
|  | Ovary | 305 | 28478 | 7631 |  |
|  | Pancreas | 166 | 17221 | 3985 |  |
|  | Prostate | 493 | 16424 | 5363 |  |
|  | Skin | 466 | 206907 | 43882 |  |
|  | Stomach | 411 | 103103 | 24497 |  |
|  | Thyroid | 488 | 4425 | 1218 |  |
|  | Uterus | 528 | 415499 | 109125 |  |
| INDEL | Bladder | 389 | 2890 | 3532 | 1.22 |
|  | Brain | 130 | 366 | 375 |  |
|  | Breast | 888 | 5207 | 6123 |  |
|  | Cervix | 257 | 3095 | 4234 |  |
|  | Colorectal | 520 | 21327 | 29876 |  |
|  | Head and Neck | 478 | 3569 | 4913 |  |
|  | Kidney | 648 | 5318 | 6395 |  |
|  | Liver | 353 | 2120 | 2143 |  |
|  | Lung | 983 | 10790 | 12407 |  |
|  | Ovary | 296 | 4260 | 4441 |  |
|  | Pancreas | 141 | 457 | 806 |  |
|  | Prostate | 416 | 1448 | 2119 |  |
|  | Skin | 430 | 1965 | 2557 |  |
|  | Stomach | 393 | 19912 | 24838 |  |
|  | Thyroid | 248 | 1746 | 1657 |  |
|  | Uterus | 516 | 27934 | 30714 |  |
| Fusion | Bladder | 269 | 809 | 819 | 0.88 |
|  | Brain | 92 | 244 | 245 |  |
|  | Breast | 787 | 3390 | 3457 |  |
|  | Cervix | 139 | 280 | 215 |  |
|  | Colorectal | 167 | 328 | 231 |  |
|  | Head and Neck | 285 | 532 | 530 |  |
|  | Kidney | 171 | 318 | 420 |  |
|  | Liver | 179 | 513 | 423 |  |
|  | Lung | 716 | 1992 | 710 |  |
|  | Ovary | 316 | 983 | 609 |  |
|  | Pancreas | 72 | 130 | 161 |  |
|  | Prostate | 381 | 1079 | 1120 |  |
|  | Skin | 287 | 841 | 918 |  |
|  | Stomach | 178 | 559 | 486 |  |
|  | Thyroid | 110 | 140 | 250 |  |
|  | Uterus | 85 | 501 | 499 |  |

### Shared neoantigens generated from frequent somatic mutations

Currently, most of the neoantigen-targeted immunotherapies are personalized and costly, which makes us wonder if we can identify frequently shared neoantigens that can be applied to a wider range of tumor patients. Here we analyzed the frequency of each neoantigen and obtained 16,913 neoantigens shared in at least two tumor samples (Table S1). Among three SNV-derived neoantigens shared in more than 20 samples, the mutated peptides are generated from *BRAF* and *KRAS*, which are well-known cancer driver genes. The most frequent shared neoantigen derived from SNV is the complex of HLA-B57:01 and mutated peptide GLAT*<u>E</u>*KSRW generated by BRAF V600E, which is present in 41 tumor samples. The complex of HLA-A02:01 and neo-peptides RLMAPVGSV, SLLTQPSPA generated by the frame-shift mutation XYLT2 G529Afs*78 are the most frequent neoantigens among INDEL-derived neoantigens, which are both appearing in 31 samples (Table 2). The two Fusion-derived shared neoantigens are the complex of HLA-A02:01 and neo-peptides ALNS*<u>EALSVV</u>* and ALNS*<u>EALSV</u>* generated by the fusion of gene *TMPRSS2* and *ERG,* which are both appearing in 14 samples (Table S1). We believe that these shared neoantigens are expected to be ideal drug targets for tumor immunotherapy, which might need further experimental validation.

**Table 2.** The detailed information of shared neoantigens occurred in more than 20 samples.

| **Mutation type** | **Mutation** | **HLA** | **peptide** | **No. of sample** |
| --- | --- | --- | --- | --- |
| SNV | BRAF_V600E_5 | HLA-B57:01 | GLATEKSRW | 41 |
| SNV | KRAS_G12D_1 | HLA-B08:01 | DGVGKSAL | 37 |
| INDEL | XYLT2_G529Afs*78 | HLA-A02:01 | RLMAPVGSV | 31 |
| INDEL | XYLT2_G529Afs*78 | HLA-A02:01 | SLLTQPSPA | 31 |
| SNV | KRAS_G12V_5 | HLA-A03:01 | VVGAVGVGK | 30 |
| INDEL | SETD1B_H8Tfs*27 | HLA-C07:01 | LRARGGTTI | 29 |
| INDEL | PLEKHA6_V328Yfs*172 | HLA-A02:01 | IMMSWMPPL | 25 |
| INDEL | PLEKHA6_V328Yfs*172 | HLA-A02:01 | VLSGCHLAV | 25 |
| INDEL | XYLT2_G529Afs*78 | HLA-C07:02 | GRTPTTRLM | 25 |
| INDEL | PLEKHA6_V328Yfs*172 | HLA-A02:01 | FTPLSAHPV | 25 |
| INDEL | PLEKHA6_V328Yfs*172 | HLA-A02:01 | CLAGSLSTM | 25 |
| INDEL | PLEKHA6_V328Yfs*172 | HLA-A02:01 | SAMPSAMGV | 25 |
| INDEL | PLEKHA6_V328Yfs*172 | HLA-A02:01 | SIMMSWMPPL | 25 |
| INDEL | XYLT2_G529Afs*78 | HLA-B07:02 | RPACTCISM | 23 |
| INDEL | XYLT2_G529Afs*78 | HLA-B07:02 | HPQWAPHSA | 23 |
| INDEL | XYLT2_G529Afs*78 | HLA-B07:02 | RPTGRTPTTRL | 23 |
| INDEL | XYLT2_G529Afs*78 | HLA-B07:02 | SPGACRPAC | 23 |
| INDEL | XYLT2_G529Afs*78 | HLA-B07:02 | RPTGRTPTT | 23 |
| INDEL | MUC4_L1937Pfs*1069 | HLA-B07:02 | HPQVTPPL | 23 |
| INDEL | MUC4_L1937Pfs*1069 | HLA-B07:02 | HPQVTPPLF | 23 |
| INDEL | SETD1B_H8Tfs*27 | HLA-C07:02 | LRARGGTTI | 22 |
| INDEL | MUC4_L1937Pfs*1069 | HLA-B07:02 | LPQHPQVTPPL | 21 |

### Experimentally validated neoantigens

In the “Validation” page of TSNAdb v1.0, we only collected experimental data about pairs of wild-type peptides and HLA alleles which hardly to be identified as neoantigens due to the limited binding data between mutated peptides and HLA alleles. With the development of clinical studies on neoantigen-based tumor immunotherapy, a large number of experimental results have provided a rich source for the functional confirmation of neoantigens. Here we collected 1,856 experimentally validated pairs of mutated peptides and HLA alleles from publications and other databases. They are divided into three tiers according to the level of experimental validation, among which 67 neoantigens are classified as tier1, 1,190 neoantigens are classified as tier2, and 599 neoantigens are classified as tier3 (see Methods in detail). Among the collected neoantigens, most of them are SNV-derived (22 are Fusion-derived, 125 are INDEL-derived, 23 are non-coding-derived, 33 are RNA splice-derived and the remains are SNV-derived) and enriched in several tumor types (430 belong to lung cancer, 477 belong to skin cancer, 361 belong to B-cell lymphoma, 123 belong to colorectal cancer).

### The usage of TSNAdb v2.0

The web interface of TSNAdb v2.0 contains seven pages: ‘Home’, ‘Browse’, ‘Search’, ‘Collected’, ‘Tools’, ‘Download’ and ‘Help’. The pages ‘Home’, ‘Download’ and ‘Help’ are similar to those presented in TSNAdb v1.0. The ‘Tools’ page is newly added to provide the links of DeepHLApan and TSNAD which we developed previously for neoantigen prediction. Major changes (such as more presentation forms, more correlation analysis, more meaningful links, and so on) have been made in the pages ‘Browse’, ‘Search’ and ‘Collected’ compared to the TSNAdb v1.0.

On the ‘Browse’ page, three subpages (‘mutation types’, ‘tumor types’, and ‘shared neoantigens’) are provided for customized neoantigen browsing. In ‘mutation types’, four parts include the ‘Statistics’ (Figure 3a), ‘Neoantigen with mutation’ (Figure 3b), ‘Neoantigen with clinical information (Figure 3c), and ‘Detailed neoantigens’ (Figure 3d) are displayed. In ‘tumor types’, three parts including the ‘Statistics’ (Figure 3e), ‘Neoantigen with clinical information (Figure 3f), and ‘Detailed neoantigen’ (Figure 3g) are displayed. In the ‘shared neoantigens’, the distribution of shared neoantigens that occur in at least two tumor samples is displayed. The table below the boxplot displays the shared neoantigens which meet different thresholds. The genes and mutations are linked to CandrisDB [15] to check whether they are driver genes or driver mutations since shared neoantigens derived from driver mutations would be potential ideal targets for tumor immunotherapy (Figure 3h).

**Figure 3.**
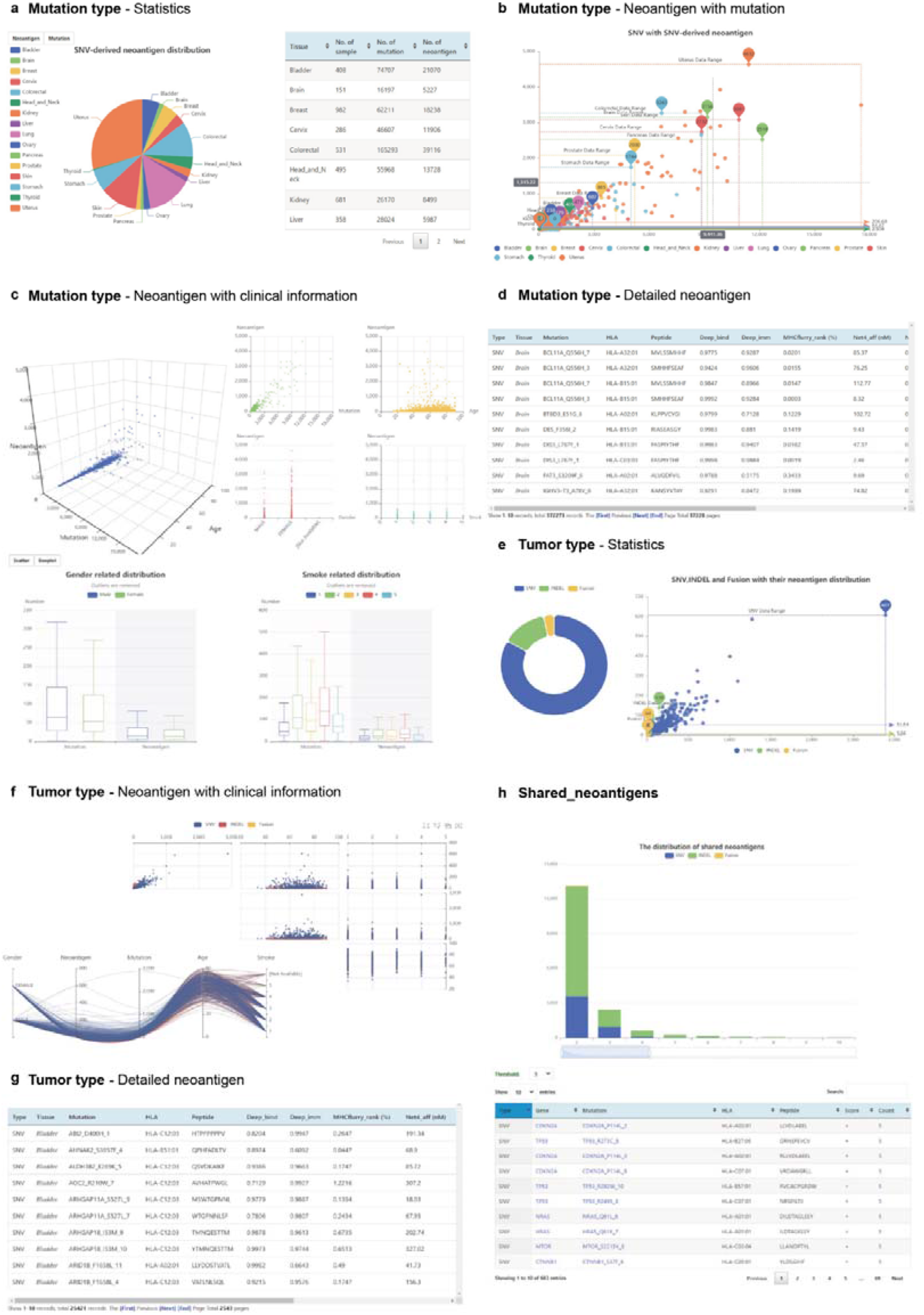
Screenshots of the Browse page of TSNAdb v2.0. **a.** The “Statistics” part in the “Mutation Type” page. **b.** The “Neoantigen with mutation” part in the “Mutation Type” page. **c.** The “Neoantigen with clinical information” part in the “Mutation Type” page. **d.** The “Detailed neoantigen” part in the “Mutation Type” page. **e.** The “Statistics” part in the “Tumor Type” page. **f.** The “Neoantigen with mutation” part in the “Tumor Type” page. **g.**The “Detailed neoantigen” part in the “Tumor Type” page. **h.** The “Shared neoantigens” page.

The ‘Search’ page contains the main page and two subpages ‘Gene’ and ‘HLA’. On the main page of ‘Search’, users could search for desired neoantigens by selecting the mutation type, tumor type, and gene. Compared with the ‘Detailed neoantigen’ of the ‘Browse’ page, it could provide more customized functions such as sorting, searching, and so on. In the subpages ‘Gene’ and ‘HLA’, the detailed neoantigens and their distribution of selected genes or HLAs would be displayed once searching. The displayed pie charts are linked with the bellowed table that the detailed neoantigens would be changed once clicking on the part of the pie charts.

On the ‘Collected’ page, all collected neoantigens are validated by experiments to be presented to the cell surface or immunogenic, which are different from those in TSNAdb v1.0. The corresponding genes and mutations of neoantigens are also linked to CandrisDB as those in the ‘shared neoantigens’.

## DISCUSSION

Neoantigens play an important role in tumor immunotherapy. A comprehensive and high confident neoantigen database would greatly meet the needs of clinical research. In TSNAdb v2.0, we predict more mutation type-derived neoantigens with stricter criteria, present the tissue-specific and gene-specific distribution of candidate tumor-specific neoantigens of TCGA tumor samples, and collect 1,557 neoantigens that have been experimentally validated, which is the most systematic database of tumor-specific neoantigens at present. Compared with other databases, TSNAdb v2.0 has several advantages as follows. Firstly, TSNAdb v2.0 provides both high-quality predicted neoantigens and experimentally validated neoantigens that most of the other databases except NEPdb only provide one of them. Compared with NEPdb, TSNAdb v2.0 provides more sources of predicted neoantigens and has richer forms of presentation. Secondly, TSNAdb v2.0 provides the analysis of shared neoantigens and links corresponding genes and mutations to CandrisDB to find out high-quality neoantigens, which other databases don’t provide. Last but not the least, TNSAdb would be updated continuously to provide constant service for related researchers and clinicians. We believe that it would certainly contribute to neoantigen-based tumor immunotherapy.

However, neoantigens are not only derived from SNVs, INDELs, and Fusions but also derived from splice variants [24], mitochondrial genome [24], and translated unannotated open reading frames [25], and so on. It’s necessary to predict all sources of neoantigen to construct a comprehensive neoantigen database. Limited by the difficulty of collecting other mutations and corresponding HLAs, we only choose three sources of neoantigen in this version of the database. In the following update of TNSAdb, we would add neoantigens from more sources and collect validated neoantigens timely to construct a more comprehensive neoantigen database.

## DATA AVAILABILITY

TSNAdb v2.0 is freely available at https://pgx.zju.edu.cn/tsnadb/.

## ACKNOWLEDGEMENTS

This work was supported by the National Natural Science Foundation of China [Grant No. 31971371, U20A20409], the Key R&D Program of Zhejiang Province [Grant No. 2020C03010], the Huadong Medicine Joint Funds of the Zhejiang Provincial Natural Science Foundation of China [Grant No. LHDMZ22H300002], the Alibaba-Zhejiang University Joint Research Center of Future Digital Healthcare. We thank the Information Technology Center, State Key Lab of CAD&CG and Innovation Institute for Artificial Intelligence in Medicine, Zhejiang University, and also Alibaba Cloud for the support of computing resources. We also gratefully acknowledge Prof. Feng Zhu from Zhejiang University for the critical reading of the manuscript, and the clinical contributors and data producers from the TCGA Research Network for referencing the TCGA datasets and the TCIA for referencing HLA-type data of TCGA samples.

